# Integrative analysis to identify shared mechanisms between schizophrenia and bipolar disorder and their comorbidities

**DOI:** 10.1101/2022.03.07.483233

**Authors:** Vinay Srinivas Bharadhwaj, Sarah Mubeen, Astghik Sargsyan, Geena Mariya Jose, Stefan Geissler, Martin Hofmann-Apitius, Daniel Domingo-Fernández, Alpha Tom Kodamullil

## Abstract

Schizophrenia and bipolar disorder are characterized by highly similar neuropsychological signatures, implying shared neurobiological mechanisms between these two disorders. These disorders also have comorbidities with other indications, such as type 2 diabetes mellitus (T2DM). To date, an understanding of the mechanisms that mediate the link between these two disorders remains incomplete. In this work, we identify and investigate shared patterns across multiple schizophrenia, bipolar disorder and T2DM gene expression datasets through multiple strategies. Firstly, we investigate dysregulation patterns at the gene-level and compare our findings against disease-specific knowledge graphs (KGs). Secondly, we analyze the concordance of co-expression patterns across datasets to identify disease-specific as well as common pathways. Thirdly, we examine enriched pathways across datasets and disorders to identify common biological mechanisms between them. Lastly, we investigate the correspondence of shared genetic variants between these two disorders and T2DM as well as the disease-specific KGs. In conclusion, our work reveals several shared candidate genes and pathways, particularly those related to the immune and nervous systems, which we propose mediate the link between schizophrenia and bipolar disorder and its shared comorbidity, T2DM.

## 1. Introduction

Psychiatric illnesses, such as schizophrenia and bipolar disorder, are known to be two of the most prevalent forms of mental disorders (Jongsma *et al*., 2019). Schizophrenia can be classified as a behavioral and cognitive syndrome, primarily characterized as having manifestations of psychosis, the onset of which is determined by genetic and/or environmental causes (Insel, 2010; Owen *et al*., 2016). Bipolar disorder is categorized as a chronic mood disorder, distinguished by frequent disruptions to mood, with shifts between either manic, depressive or mixed states (Grande *et al*., 2016; Müller-Oerlinghausen *et al*., 2002). While both disorders point to some degree of cognitive impairment, as of yet, there are no neuropsychological signatures that can conclusively lead to a differential diagnosis between these two conditions (Bortolato *et al*., 2015), hinting at the substantial overlap between them. These disorders have also been linked to several comorbidities, mainly type 2 diabetes mellitus (T2DM) and cardiovascular disorders (Mizuki *et al*., 2021). For instance, T2DM has been reported to have a higher prevalence in patients with schizophrenia (Mukherjee *et al*., 1989; Argo *et al*., 2011), while relatives of patients with schizophrenia have also been shown to have a higher risk of diabetes (Miller *et al*., 2016). T2DM has also been linked to bipolar disorder, as shown by a genetic analysis conducted by Torkamani *et al*. (2008), where numerous top-ranking SNPs were found in common between bipolar disorder and T2DM.

Transcriptomic data are commonly used in the psychiatric domain to gain insights on the neurobiological underpinnings of mental disorders (Perez *et al*., 2021; Song *et al*., 2021; Chaumette *et al*., 2019). Additionally, Genome Wide Association Studies (GWAS) have also shed light on the genetic variants associated with psychiatric conditions, including schizophrenia and bipolar disorder (Mullins *et al*., 2021; Smeland *et al*., 2020; Psychiatric Genomics Consortium, 2018), as well as schizophrenia associated copy number variants (Ehrhart *et al*., 2022), both of which assist in understanding the molecular mechanisms of these conditions. Nonetheless, the analysis of large-scale datasets can be challenging, prompting the construction of networks to integrate and organize these datasets into models describing biological relations which can aid in identifying gene expression patterns as well as potential disease-relevant genes (Parikshak *et al*., 2015). In the psychiatric field, gene co-expression networks have been used to identify candidate genes and pathways with potential roles in the pathobiology of bipolar disorder (Liu *et al*., 2019; Zhang *et al*., 2021). Similarly, co-expression network analysis of transcriptomic data has identified cognitive abnormalities in schizophrenic patients, revealing that variations observed in gene expression patterns across cortical regions of controls are attenuated in persons with schizophrenia (Roussos *et al*., 2012), while other studies have identified disease modules associated with this disorder (de Jong *et al*., 2012; Torkamani *et al*., 2010).

Another major technique to guide the interpretation of transcriptomic data is through the use of prior knowledge, such as in the form of pathways or knowledge graphs, to enable the contextualization of signals from omics data analyses. For instance, enrichment analysis can be used on clusters within co-expression networks (i.e., modules) to correlate modules to particular functions or biological pathways. To date, several pathways implicated in both schizophrenia and bipolar disorder have been identified, including immune dysfunction (Goldsmith *et al*., 2016) and Wnt signaling (Hoseth *et al*., 2018). In a recent joint analysis of schizophrenia and bipolar disorder, functional enrichment of modules from co-expression networks of these two disorders revealed processes which were both unique to and shared across these conditions, also implicating the immune system (Sahu *et al*., 2020). Another recent approach also jointly investigated these disorders through the identification of common pathways using a system biology approach, revealing those associated with the stress response, energy systems and neuron systems. (Altaf-Ul-Amin *et al*., 2021). Thus, due to the polygenic nature of these overlapping disorders, analyses at not only the genetic level but also the pathway and network level are crucial to elucidating their complex pathophysiology.

In this work, we aim at identifying shared mechanisms between schizophrenia and bipolar disorder at the genetic and pathway levels using an integrative approach that leverages prior knowledge and experimental data. To achieve this goal, we first identify pathways that are linked to schizophrenia and bipolar disorder through three major approaches, namely, a meta-analysis, a co-expression network analysis, and pathway enrichment. In doing so, we attempt to reveal the common mechanisms that may mediate the link between schizophrenia and bipolar disorder. Finally, we emphasize the critical importance of an integrative, data- and knowledge-driven approach in order to better understand the comorbid associations of psychiatric disorders.

## 2. Methodology

In subsection 2.1, we outline the process of acquiring datasets for the two psychiatric disorders. In subsection 2.2, we detail the generation of a knowledge graph (KG) for schizophrenia and bipolar disorder. Then, in subsection 2.3, we describe the procedure to perform a meta-analysis on the datasets **(Figure 1A(i))**, while in 2.4, the process of generating co-expression networks from transcriptomic datasets is outlined **(Figure 1A(ii))**. Next, in subsection 2.5, we describe the pathway enrichment analyses performed **(Figure 1A(iii))**. Finally, in section 2.6, we outline an equivalent comorbidity analysis conducted for T2DM.

**Figure 1.**
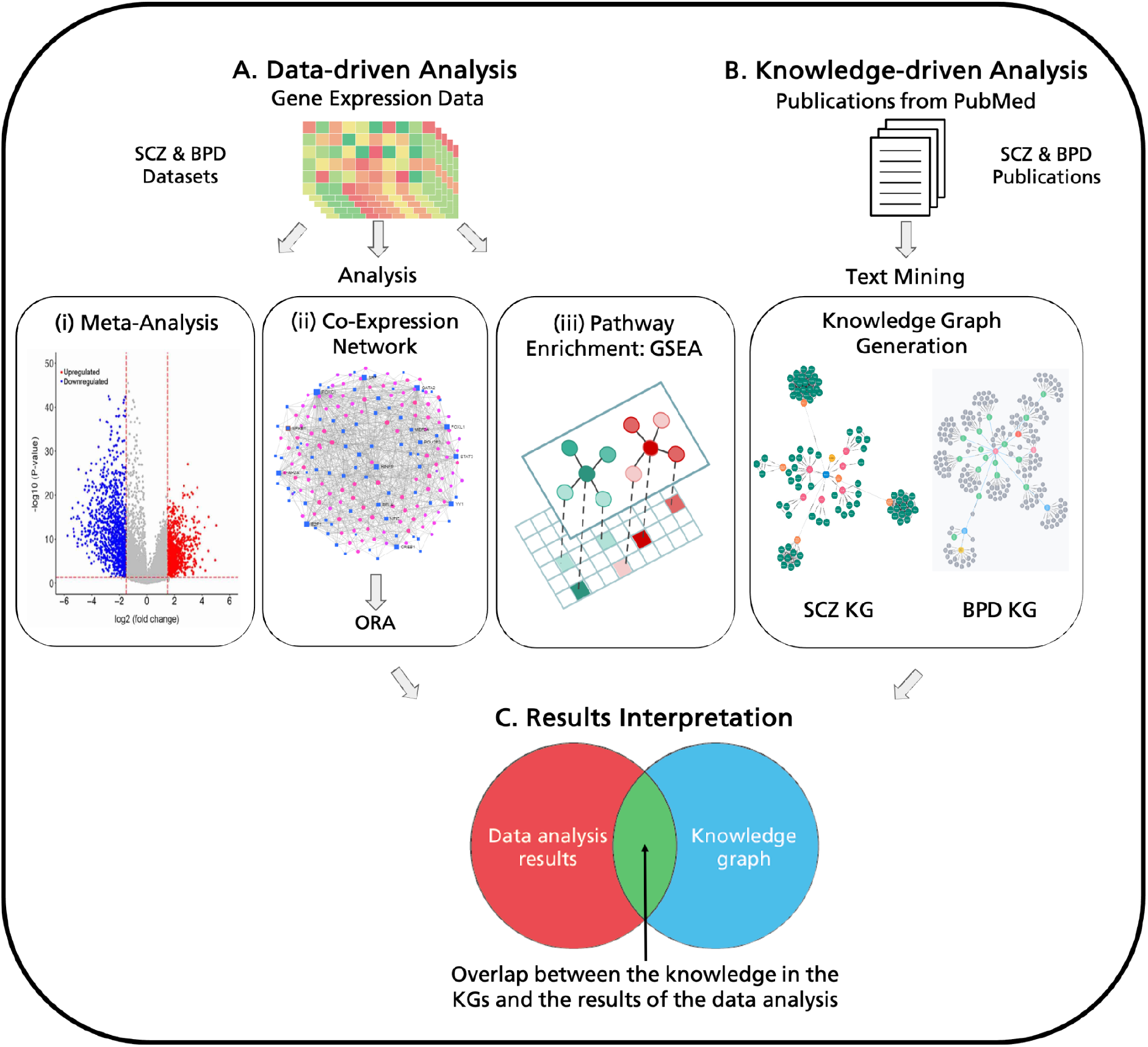
Workflow for the methodology of the paper. The methodology is divided into two parts, **A)** gene expression data analysis and **B)** prior knowledge integration. **A)** Gene expression datasets for schizophrenia (SCZ) and bipolar disorder (BPD) are collected and analyzed through three different approaches: (i) meta-analysis of the gene expression datasets, (ii) co-expression network analysis in conjunction with pathway enrichment using over-representation analysis (ORA), and (iii) pathway enrichment with gene set enrichment analysis (GSEA). **B)** Text mining is performed on publications available in PubMed to generate disease-specific knowledge graphs (KGs). **C)** Results from the data analysis and knowledge curation are compared to identify the overlap between these parallel approaches.

### 2.1. Collection of genetic and gene expression datasets

In order to carry out the data-driven analysis **(Figure 1A)**, we collected gene expression datasets from schizophrenia and bipolar disorder. Microarray datasets for schizophrenia and bipolar disorder were collected from ArrayExpress and Gene Expression Omnibus (GEO) on March 2nd, 2021. Our selection criteria for the datasets were two-fold: i) datasets must have been generated using the Affymetrix GeneChip Human Genome U133 family platform from human patient samples to avoid platform variability, and ii) datasets must have contained more than 40 control and disease samples. The datasets that fulfilled this criteria for schizophrenia included E-GEOD-12649, E-GEOD-21138, E-GEOD-21935, E-GEOD-53987, and GSE93987 **(Supplementary Table 1)**. Datasets fulfilling this criteria for bipolar disorder included E-GEOD-46449, E-GEOD-5388, E-GEOD-5392, E-GEOD-53987, and GSE12649 **(Supplementary Table 2)**. Samples for the datasets were taken from tissues in the brain, except for samples from the bipolar disorder dataset E-GEOD-46449, which contained samples taken from leukocytes in the blood (see **Supplementary Tables 1 and 2)**.

The schizophrenia and bipolar disorder datasets were pre-processed using RMA normalization with the R package, oligo (Carvalho and Irizarry, 2010). Gene probes of the datasets were also annotated to HGNC gene symbols with the help of the annotateEset function from affycoretools (MacDonald, 2020) and the annotation data libraries from affymetrix.

Lastly, we leveraged single nucleotide polymorphisms (SNPs) associated with schizophrenia and bipolar disorder (*p*-value < 5.0 x 10^-8^) from GWAS Catalog (downloaded on 11.02.2022) (Buniello *et al*., 2019). A total of 2,425 and 1,168 SNPs for schizophrenia and bipolar disorder were respectively mapped to 1,610 and 984 genes of which 265 genes were shared (https://github.com/vinaysb/psychiatric_disorders_corpus).

### 2.2. Disease-specific knowledge graphs

In order to represent known interactions from the literature around each disease, we built disease-specific KGs for schizophrenia and bipolar disorder. The majority of the KGs were automatically generated by converting the text in publications into Biological Expression Language (BEL) statements with the help of a proprietary text-mining workflow (Geißler, 2020). Automatic text mining was carried out on abstracts of publications whereas manual curation was done on full texts. Consequently, KGs generated from text-mining were merged with KGs generated from manual curation (https://github.com/vinaysb/psychiatric_disorders_corpus). The articles were retrieved from PubMed using advanced search with the combined MeSH terms “Schizophrenia” and “Bipolar Disorder” for scientific articles published in the last 10 years. The search resulted in approximately 90,000 articles which were then converted into a computer-readable network in the form of BEL statements. The BEL files of the schizophrenia (SCZ) KG contained 2,380 unique nodes and 19,984 edges **(Supplementary Figures 1 and 2)**, while the bipolar disorder (BPD) KG contained 2,042 unique nodes and 16,350 edges **(Supplementary Figures 3 and 4)**.

### 2.3. Meta-analysis of gene expression datasets for schizophrenia and bipolar disorder

In order to identify which genes were up- and down-regulated across all datasets of a particular disease, a meta-analysis was carried out on the schizophrenia and bipolar datasets separately. Subsequently, genes that were consistently dysregulated across all datasets were identified and compared against each other.

Prior to conducting the meta-analysis, each of the pre-processed datasets were further processed through the limma R package (Ritchie *et al*., 2015) to calculate the differential expression of each gene in the disease state versus controls. Differential expression was calculated via log_2_ fold changes (log_2_FC), which were corrected using the Benjamini-Hochberg (BH) multiple test corrections method (Benjamini and Hochberg, 1995). Differentially expressed genes for each dataset were subsequently filtered to include only those with an adjusted *p*-value < 0.05, which were then used to carry out the meta-analysis and identify genes that were consistently over- or under-expressed. The meta-analysis was conducted using a vote-counting technique which counts the number of positive, negative and non-significant changes in a dataset (Bushman and Wang, 2009) implemented in the MetaVolcanoR (Prada *et al*., 2020) package.

### 2.4. Generating co-expression networks

Co-expression network analysis was performed to identify correlation patterns between transcripts measured in the gene expression datasets. Co-expression networks were generated for the disease samples in each dataset with the WGCNA package in R (Langfelder *et al*., 2008). The networks were subsequently used to identify correlations that were common between the disease samples of all the datasets for schizophrenia and bipolar disorder. To create the co-expression networks, the WGCNA package uses an unsigned similarity measure to generate a topological overlap matrix (TOM) based on the similarity of their expression pattern. The TOM was then pruned to retain the top 1% highest similarity in it to build the co-expression network. The individual networks constructed for each of the datasets were then used to identify common edges across all datasets for schizophrenia and bipolar disorder.

### 2.5. Pathway enrichment analysis

To determine the pathways affected by dysregulated genes in the datasets, we employed two different pathway enrichment methods (i,e., over-representation analysis (ORA) and Gene Set Enrichment Analysis (GSEA) (Subramanian *et al*., 2005)) and 337 pathways from the KEGG database (retrieved 3rd August 2020, version 95.2) (Kanehisa, *et al*., 2021) using DecoPath (Mubeen *et al*., 2021). ORA uses the hypergeometric test to assess whether a pathway is significantly enriched in a list of genes. We conducted ORA on the gene list from common edges of the co-expression networks, while GSEA was performed on processed gene expression datasets using the GSEApy library (Fang, 2020). GSEA calculates the enrichment score employing a Kolmogorov–Smirnov-like statistic before estimating the statistical significance by calculating *p*-values for the results which are corrected *via* the Benjamini–Hochberg method (Benjamini and Hochberg, 1995). Finally, the enriched pathways of the datasets of both diseases were checked for overlap.

### 2.6. Comorbidity analysis

In order to identify common mechanisms between schizophrenia and bipolar disorder which could explain their shared comorbidity with T2DM, we also conducted the above-mentioned analyses (see **Sections 2.3 – 2.5**) on T2DM datasets. We leveraged T2DM datasets from Karki *et al*. (2017) in which the authors investigated a comorbid association between Alzheimer’s disease and T2DM, identifying four datasets which fulfilled the criteria outlined in **Section 2.1** (see **Supplementary Table 3**). Furthermore, we used a T2DM KG that was previously manually curated by Karki *et al*. (2020). Finally, we leveraged SNPs associated with T2DM (p-value < 5.0 x 10-8) from GWAS Catalog (downloaded on 25.02.2022) (Buniello et al., 2019). A total of 3,172 SNPs were mapped to 2,026 genes.

## 3. Results

In subsection 3.1, we outline the results of a meta-analysis using several gene expression datasets for schizophrenia, bipolar disorder and T2DM. In subsection 3.2, we utilize these datasets to generate co-expression networks for the two psychiatric disorders as well as T2DM and investigate the overlap across these networks. Finally, in subsection 3.3, we carried out pathway enrichment analysis on the gene expression datasets for these three disorders and identified the shared mechanisms and overlap of enriched pathways with the KGs.

### 3.1. Comparing consistently differentially expressed genes against literature

A meta-analysis was conducted in order to identify genes that were dysregulated across datasets for schizophrenia and bipolar disorder. The results of the analysis showed that the majority of dysregulated genes were either up- or down-regulated in a single dataset and only a small minority were dysregulated in more than one dataset **(Supplementary Figure 5)**. From this meta-analysis, we obtained the consensus results on the direction of dysregulation for genes from these two disorders **(Table 1)**. We observed 15 genes that were consistently dysregulated in the schizophrenia datasets and 37 in the bipolar disorder datasets. For T2DM, we relied upon the results from Karki *et al*., (2017). Here, we found that none of the genes were either significantly up- or down-regulated across all datasets (p-value < 0.05).

**Table 1.** List of genes that were consistently up- or down-regulated in two or more gene expression datasets for **(a)** schizophrenia and **(b)** bipolar disorder.

| a) Schizophrenia |  | b) Bipolar Disorder |  |  |  |
| --- | --- | --- | --- | --- | --- |
| Gene | Regulation | Gene | Regulation | Gene | Regulation |
| ACTR3B | Down-regulated | AASDHPPT | Down-regulated | SKP1 | Down-regulated |
| ATP2A2 | Down-regulated | ARPP19 | Down-regulated | SLC35D1 | Down-regulated |
| CLK4 | Down-regulated | BCL11A | Down-regulated | SRRM1 | Down-regulated |
| COPG2 | Down-regulated | CAPZA2 | Down-regulated | STRN3 | Down-regulated |
| DCTN1 | Down-regulated | FABP5 | Down-regulated | SUB1 | Down-regulated |
| ENSA | Down-regulated | HNRNPDL | Down-regulated | TLK1 | Down-regulated |
| HNRNPA3 | Down-regulated | HNRNPH1 | Down-regulated | TMED2 | Down-regulated |
| NDUFA10 | Down-regulated | HNRNPU | Down-regulated | UBE3A | Down-regulated |
| NDUFS1 | Down-regulated | IMPACT | Down-regulated | UFM1 | Down-regulated |
| PDAP1 | Down-regulated | MEAF6 | Down-regulated | WBP11 | Down-regulated |
| SFSWAP | Down-regulated | IPO5 | Down-regulated | CLASRP | Up-regulated |
| YWHAZ | Down-regulated | MMD | Down-regulated | LIPE | Up-regulated |
| SAT1 | Up-regulated | MRPS33 | Down-regulated | LRRFIP1 | Up-regulated |
| SLC16A6 | Up-regulated | MTX2 | Down-regulated | NEK11 | Up-regulated |
| ZEB1 | Up-regulated | PNISR | Down-regulated | NTRK3 | Up-regulated |
|  |  | PTS | Down-regulated | RAB35 | Up-regulated |
|  |  | RAB40B | Down-regulated | SLC6A8 | Up-regulated |
|  |  | RAP2B | Down-regulated | TCP10L | Up-regulated |
|  |  | SFPQ | Down-regulated |  |  |

In order to identify the overlap between the results of the meta-analysis on the schizophrenia dataset and the SCZ KG, we compared the list of consistently up- or down-regulated genes **(Table 1)** with the nodes of the KG. We were able to find some concordance between the results of the meta-analysis and the KG, through which we could explore known interactions of the affected genes. For instance, DCTN1 was found to be consistently down-regulated across datasets and was identified in the SCZ KG. In contrast, this gene has been shown to exhibit increased expression in patients with psychiatric disorders and is considered a blood biomarker for the detection of these disorders (Kurian *et al*., 2011). Furthermore, Yu *et al*. (2021) have shown a link between the gene and schizophrenia as well as psychosis, and found cross-talk between DCTN1 and STX1A. We then investigated this edge in the SCZ KG, finding a negative correlation edge between STX1A and STXBP1. Additionally, we were able to identify a positive correlation between schizophrenia and the STXBP1 gene in the KG, as reported in studies by Ramos-Miguel *et al*. (2015) and Urigüen *et al*. (2013). Consistently dysregulated genes that appeared in the KG also included NDUFS1 (Zhu *et al*., 2015), YWHAZ (Zhu *et al*., 2021), and HNRNPA3 (Mohammadi *et al*., 2018).

Similarly, we conducted an equivalent investigation for bipolar disorder. The meta-analysis revealed that NTRK3 was consistently up-regulated in several datasets. NTRK3 encodes neurotrophic tyrosine receptor kinase 3 receptor protein which is responsible for the binding of neurotrophin, which plays a key role in the MAPK signaling pathway (Reichardt, 2006). Upon examination of this gene in the BPD KG, we found an association between NTRK3 and mental disorders, based upon evidence found in Braskie *et al*., (2013). We then investigated the neighbors of NTRK3 in the KG, finding NTF3 increases NTRK3 and NTF3 is also positively correlated with bipolar disorder. Furthermore, Athanasiu *et al*. (2011) conducted a SNP-disease association study on bipolar disorder, where they identified several polymorphisms of NTRK3 with links to bipolar disorder. These polymorphisms were also closely associated with schizophrenia and corroborated in a study by Gratacòs *et al*., (2001). Finally, the KG also revealed a positive correlation between bipolar disorder and UBE3A (You *et al*., 2020), a consistently down-regulated gene in the meta-analysis.

In conclusion, using the meta-analysis we were able to identify consistently dysregulated genes for schizophrenia and bipolar disorder. However, the meta-analysis for T2DM datasets did not reveal any significantly dysregulated genes, thus we were unable to investigate genes potentially implicated in comorbid associations of the two psychiatric disorders and T2DM. While some of the genes from the meta-analysis on schizophrenia and bipolar disorder were present in the disease-specific KGs, many of them were absent from it and appear to be under studied, warranting further investigation. Nevertheless, we have shown here how the consolidation of literature knowledge in the form of KGs can guide the interpretation of gene expression data.

### 3.2. Identifying enriched pathways from co-expression patterns

In this section, we leveraged the co-expression networks generated for individual datasets to study the activity and correlation patterns of genes in both disorders. This method of analysis has already been successfully used to identify correlated genes in RNA-seq data for a single schizophrenia and bipolar disorder dataset (Sahu *et al*., 2020). Here, however, we focused on identifying the common mechanisms among schizophrenia and bipolar disorder by leveraging multiple datasets for each of the disorders. We were able to find common edges between 10 co-expression networks of the schizophrenia and bipolar disorder datasets **(Figure 2)**. To identify shared mechanisms across these diseases, we performed ORA on the gene nodes of common edges of the co-expression networks (i.e., edges that appear across all datasets for schizophrenia and bipolar disorder).

**Figure 2.**
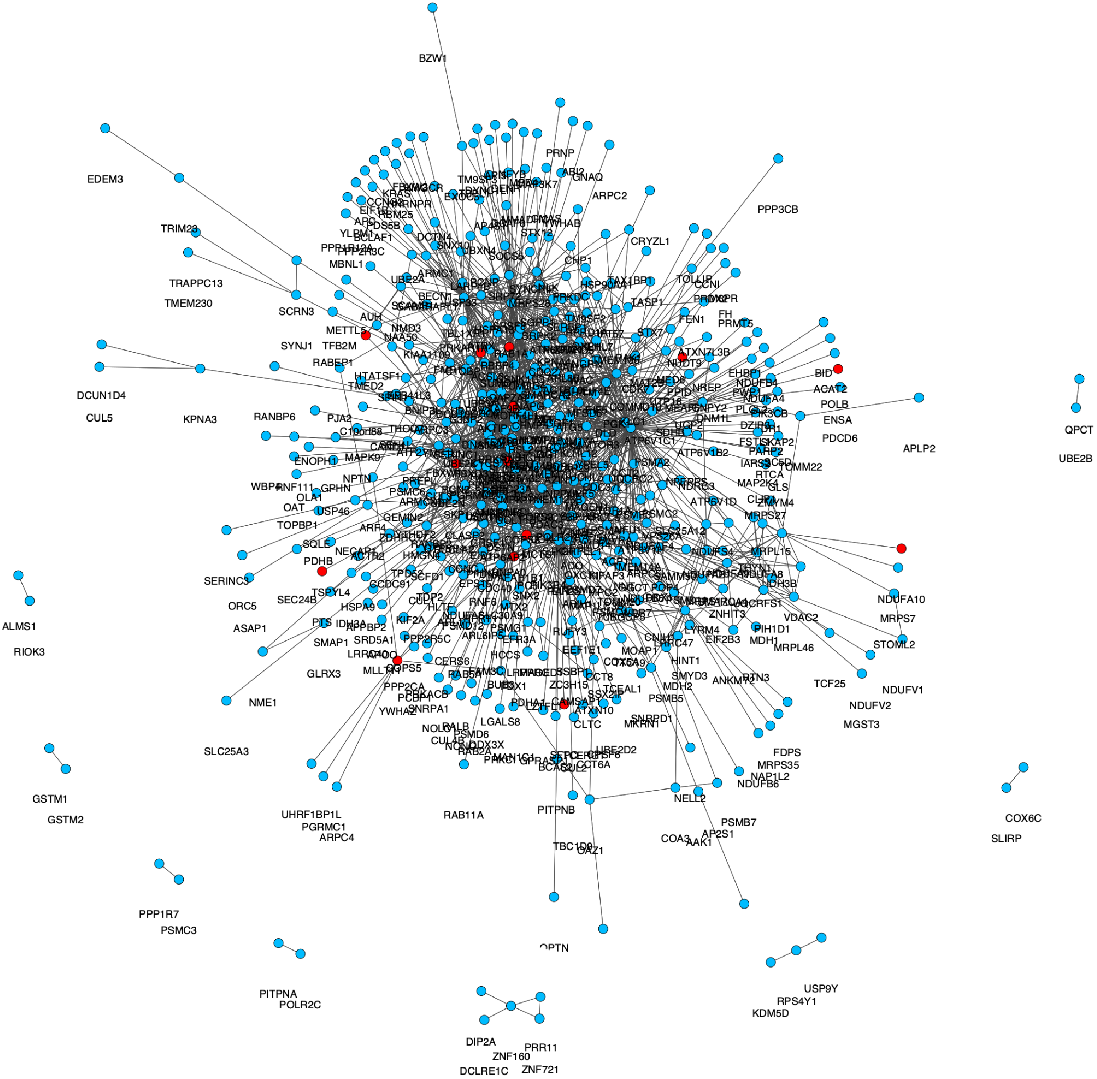
Shared edges across co-expression networks from the schizophrenia and bipolar disorder datasets. Consistently dysregulated genes from the meta-analysis of both schizophrenia and bipolar disorder are marked in red.

On average, co-expression networks corresponding to each of the schizophrenia and bipolar disorder datasets (i.e., 5 for schizophrenia and 5 for bipolar disorder) contained 2-7 million edges. We first identified the edges which occurred consistently across all datasets from each disorder, finding 16,748 and 9,550 edges in common for schizophrenia and bipolar disease, respectively.

The intersection of these disease-specific edges revealed that 1,094 edges of them were shared. We then investigated the 424 unique genes present in these edges by running ORA against 337 KEGG pathways. The results yielded 111 significantly enriched pathways (*q*-value < 0.01) (**Supplementary Table 4**), the top three ranked pathways were Hungtinton’s, Alzheimer’s, and Parkinson’s disease, all of which have a comorbidity with both schizophrenia and bipolar disorder (Rocha *et al*., 2018; Cai and Huang, 2018; Faustino *et al*., 2020). Additionally, oxidative phosphorylation was the top ranked non-disease pathway, shown to be positively correlated with bipolar disorder (Morris *et al*., 2017). Furthermore, an increase in lithium, often prescribed as a treatment for schizophrenia, was reported to increase the activity of the oxidative phosphorylation pathway (Leucht *et al*., 2015). Finally, among the enriched pathways were those which overlapped with the ones identified by Sahu *et al*. (2020) (e.g, apoptosis, autophagy and hippo signaling pathway).

The T2DM co-expression networks contained 3,389 nodes in common with the schizophrenia and bipolar disorder co-expression networks. Although we were able to identify 633 edges in common across all co-expression networks generated from the T2DM datasets, we were unable to find any common edges between the T2DM and schizophrenia networks or the T2DM and bipolar disorder networks. Nevertheless, from the 540 unique genes present in the shared edges of the T2DM co-expression networks, we were able to identify 66 enriched pathways (*q*-value < 0.05) in the ORA results (**Supplementary Table 5**). These included Hungtinton’s, Alzheimer’s and Parkinson’s disease pathways, which were also enriched in the schizophrenia and bipolar disorder ORA results. Additionally, the PI3K-Akt signaling pathway, cytokine-cytokine receptor interaction pathway, chemokine signaling pathway, and Phagosome pathway were also enriched.

### 3.3. Comparing consistently enriched pathways against pathways over-represented in the literature

We conducted pathway enrichment using GSEA on each gene expression dataset and compared the overlap of the results. For individual schizophrenia datasets, the number of significantly enriched pathways ranged from 60 to 90, while this number ranged from 40 to 50 for bipolar disorder datasets and 60 to 90 for T2DM datasets as well. For each disorder, we investigated whether any pathways were significantly enriched (*q*-value < 0.05) across all datasets, identifying four pathways across all schizophrenia datasets, three across all bipolar disorder datasets and five across all T2DM datasets (see **Supplementary Table 6** for details). To elucidate the correspondence of significantly enriched pathways detected from the data-driven analysis with the literature, these GSEA results were then compared to the results of ORA on genes from each of the three disease-specific KGs (**Supplementary Tables 7, 8 and 9)**.

Upon comparing the GSEA results of schizophrenia datasets with the SCZ KG-derived ORA results, we found that each of the enriched pathways in schizophrenia datasets were also enriched in the SCZ KG. Three of the four enriched pathways were related to immune response, including cytokine-cytokine receptor interaction, the Il-17 signaling pathway and the NF-kappa B signaling pathway, known to be related to multiple neurological disorders and schizophrenia (Azodi and Jacobson, 2016; Reale *et al*., 2021; Fang *et al*., 2018; Murphy *et al*., 2021). Through this investigation, we were also able to confirm some of the results from the previous analyses. For instance, we found that the PI3K-Akt signaling pathway was enriched in not only the pathway enrichment results of the schizophrenia datasets and the SCZ KG, but also the results of the co-expression network analysis and meta-analysis for schizophrenia.

The comparison of the GSEA results of bipolar disorder datasets with the BPD KG-derived ORA results revealed that two of the three pathways enriched in bipolar disorder datasets were also enriched in the BPD KG. These included the phagosome pathway, involved in the process of phagocytosis and linked with bipolar disorder in work carried out by Barbosa *et al*. (2015), and the antigen processing and presentation pathway, also part of the phagocytosis pathway.

When the pathway enrichment results of the T2DM datasets and T2DM KG were compared, we found minimal overlap between them. Nonetheless, the cytokine-cytokine receptor interaction pathway was enriched in both schizophrenia and T2DM. In order to further investigate shared mechanisms between the psychiatric disorders and their comorbidity, we analyzed the overlap between ORA results of the KGs for schizophrenia and T2DM (**Figure 3a**) and bipolar disorder and T2DM (**Figure 3b**). Here, we found that the vast majority of pathways enriched for T2DM were also enriched for schizophrenia and bipolar disorder.

**Figure 3.**
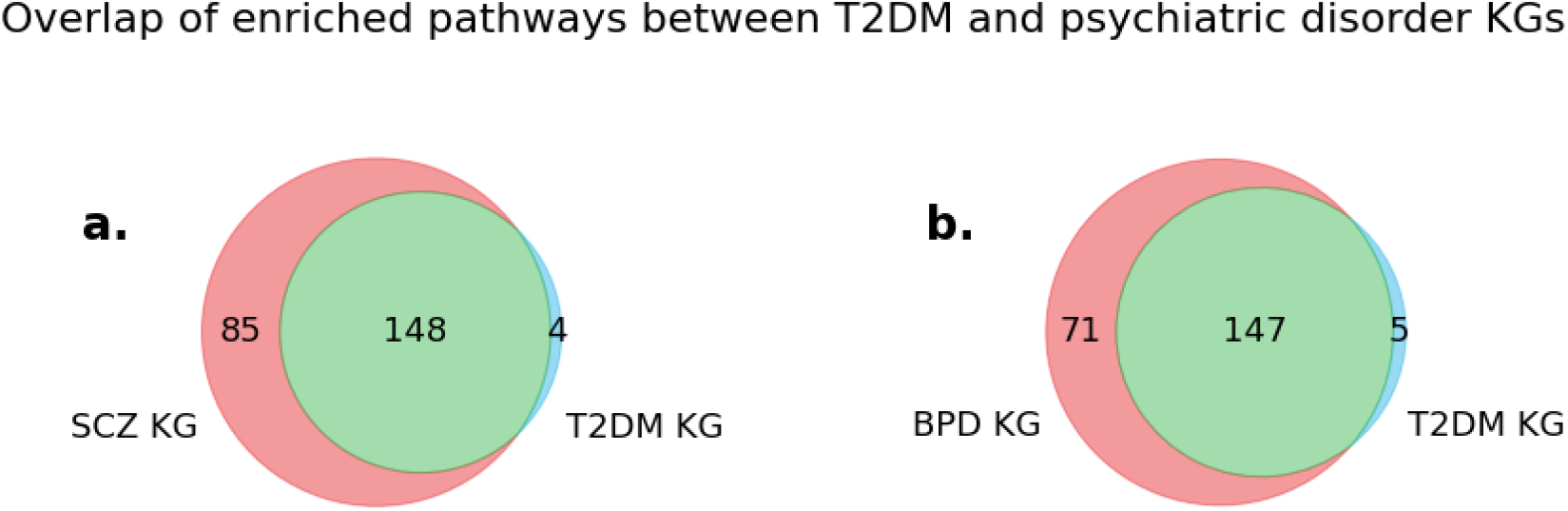
Overlap of enriched pathways between psychiatric disorders and T2DM. Venn diagram depicting the overlap between enriched pathways for the **(a)** schizophrenia and T2DM KGs and **(b)** bipolar disorder and T2DM KGs.

Next, we investigated pathways which consistently appeared across each data-driven analysis and were enriched in the disease-specific KGs. In doing so, we found several pathways, including the MAPK signaling pathway in the KG-derived ORA results, the co-expression analysis results and the meta-analysis results of schizophrenia datasets. Similarly, the neurotrophin signaling pathway was also enriched in the KG ORA results and the co-expression network analysis, while NTRK3 and NTF, both of which are involved in this pathway, were also differentially expressed in the meta-analysis on bipolar disorder datasets. Other enriched pathways such as apoptosis, hippo signaling pathway and FoxO signaling pathway, were also observed in the co-expression network analysis as well as the publication by Sahu *et al*. (2020). Additional enriched pathways such as Huntington’s, Alzheimer’s and Parkinson’s disease were not only the top three enriched pathways in the co-expression network analysis, but genes of the Huntington’s disease pathway, such as DCTN1 and STX1A, were also identified as key genes in the meta-analysis of the schizophrenia datasets. Our results are also in line with recent work by Ehrhart *et al*. (2022) who identified several enriched pathways from the WikiPathways database (Martens *et al*., 2021) from ORA on multiple gene lists of schizophrenia risk genes. More specifically, using mappings between the KEGG and WikiPathways database (Domingo-Fernández *et al*., 2018), we were able to identify an overlap of multiple pathways such as NF-kappa B signaling pathway, Huntington’s disease, T cell receptor signaling pathway and dopaminergic synapse. Furthermore, the ubiquitin pathway, which is involved in the Parkinson’s disease pathway, contains the ubiquitin ligase UBE3A, identified as a key gene in the meta-analysis of bipolar disorder. Interestingly, STX1A also appears in the seminal vesicle cycle pathway, also enriched in both the KG ORA analysis and the co-expression network analysis. Apart from the immunological pathways previously mentioned, the KG-derived ORA results also contained the TNF signaling pathway, chemokine signaling pathway, and T cell receptor signaling pathway. Pathways related to neurological disorders, such as dopaminergic synapse and GABAergic synapse, were also significantly enriched, not only in the psychiatric disorder KG ORA results, but also in the T2DM KG ORA results. Finally, we were also able to identify shared enriched immunological pathways between the results of ORA on the T2DM KG and the psychiatric disorder KGs, such as TNF signaling pathway, IL-17 signaling pathway and NF-kappa B signaling pathway.

Lastly, several studies have attempted to identify shared genetic variants as markers in both schizophrenia and bipolar disorder (Psychiatric Genomics Consortium, 2018). Thus, we investigated whether the common genetic variants for both disorders were among the genes and pathways we identified in our study. To do so, we first collected genetically associated variants for each of the diseases from the GWAS catalog (see Methods). As expected, we found that the overlap between the genetically associated variants for both indications were statistically significant (Fisher’s test; *p*-value < 0.01). Among the 265 genes shared between the two disorders, 46 and 42 were respectively present in the SCZ and BPD KGs. Interestingly, the vast majority of these latter two sets of genes present in one disease-specific KG, could also be found in the other **(Supplementary Figure 6)**. In other words, nearly all of the genes derived from disease associated genetic variants were common in the disease-specific KGs, highlighting the substantial overlap of these diseases in the literature. Notably, among these genes, we identified the above-mentioned NTRK3 in both KGs and the meta-analysis results of the bipolar disorder datasets. Similarly, ATP2A2, involved in multiple pathways including the Alzheimer’s disease pathway, calcium signaling pathway, cGMP-PKG signaling pathway and cAMP signaling pathway, was found in the results of ORA on genes present in both KGs as well as the meta-analysis results of the schizophrenia datasets. In a publication by Nakamura *et al*. (2016), the authors identified a mutation in ATP2A2 as one of the key causes for the onset of psychoses in patients with Darier’s disease. Moreover, Nakajima *et al*. (2021) further noted that mutations in ATP2A2 lead to a higher risk of developing psychiatric disorders, such as schizophrenia and bipolar disorder.

In order to further develop comorbid associations of schizophrenia and bipolar disorder with T2DM, we compared the 265 shared genes between the two psychiatric disorders with the SNPs associated with T2DM. In doing so, we were able to identify 19 genes that were shared between all three disorders (https://github.com/vinaysb/psychiatric_disorders_corpus). One notable gene out of the 19 is MSRA, whose mutation has implicated its involvement as a key candidate gene for schizophrenia (Walss-Bass *et al*., 2009) and for which a SNP has been associated with an increased risk of bipolar disorder (Ni *et al*., 2015). Additionally, in work carried out by Styskal *et al*. (2013), it was shown that MSRA knockout mice displayed insulin resistance. FYN was also found in the result of the GWAS analysis and multiple studies involving FYN knockout mice have shown the development of insulin resistance and prediabetic neuropathy (Suo *et al*., 2016; Lee *et al*., 2013). Mutated alleles of FYN have also been linked to the early onset of bipolar disorder (Szczepankiewicz *et al*., 2009)

## 4. Discussion

Here, we have presented three data-driven approaches, leveraged by prior, literature-derived and pathway knowledge, to identify candidate genes and pathways for schizophrenia and bipolar disorder, which often present with highly similar neuropsychological signatures. As both disorders have a shared comorbidity with T2DM, we conducted an equivalent approach on gene expression data from this disease to discern whether common biological mechanisms may mediate the link between T2DM and the two psychiatric disorders.

One of the main limitations of our analysis is that the T2DM datasets did not have a large variety of samples in the datasets. These datasets contain 30 to 40 samples each as compared to 70 to 80 samples per dataset for the two psychiatric disorders. Furthermore, the low number of overlapping pathways across datasets of the same disease could indicate that transcriptomic data alone may not be sufficient to reveal dysregulated patterns beyond a gene level resolution. However, a few pathways were enriched across all different analyses conducted, some of them shared (e.g., cytokine-cytokine receptor interaction), indicating that these pathways may be involved in the pathophysiology of the studied indications. Another limitation of our work is that we have solely relied upon the KEGG database. Although KEGG is a highly cited pathway resource, pathways contained in this resource may not adequately capture mechanisms within the psychiatric domain. In the future, we foresee using other modalities (e.g., proteomics) or samples from other cells (e.g., blood samples) in place of tissue samples as recent studies have shown that blood biomarkers can be effectively used to detect schizophrenia and bipolar disorder (Wagh *et al*., 2021; Munkholm *et al*., 2015).

## Authors’ Contributions

ATK and DDF conceived and designed the study. VS analyzed the datasets and interpreted the results. AS and GMJ curated the disease-specific KGs. ATK and MHA acquired the funding. VS, SM, DDF and ATK wrote the manuscript. MHA reviewed the manuscript.

All authors have read and approved the final manuscript.

