## Supplementary Figures for "Integrative analysis to identify shared mechanisms between schizophrenia and bipolar disorder and their comorbidities"

### Additional File

#### Supplementary Figures

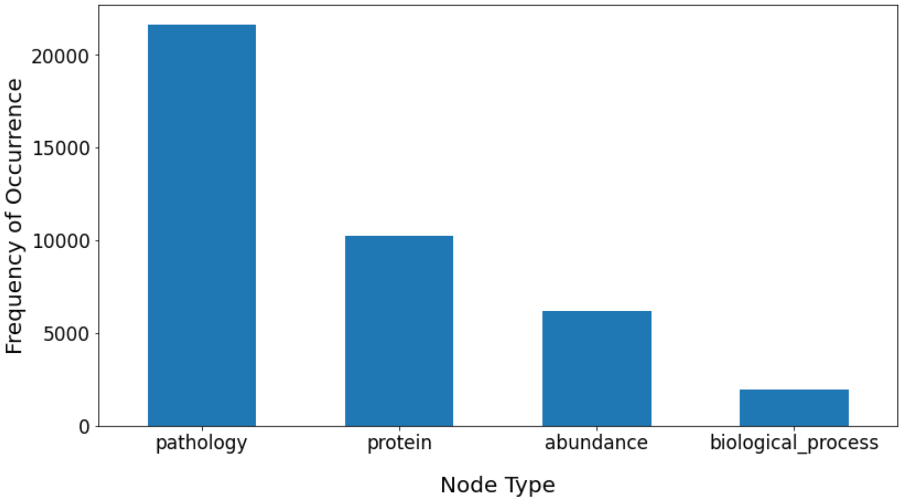


**Supplementary Figure 1. Node type statistics of the schizophrenia knowledge graph.** Bar plot representing the number of BEL statements that contain the different node/entity types.

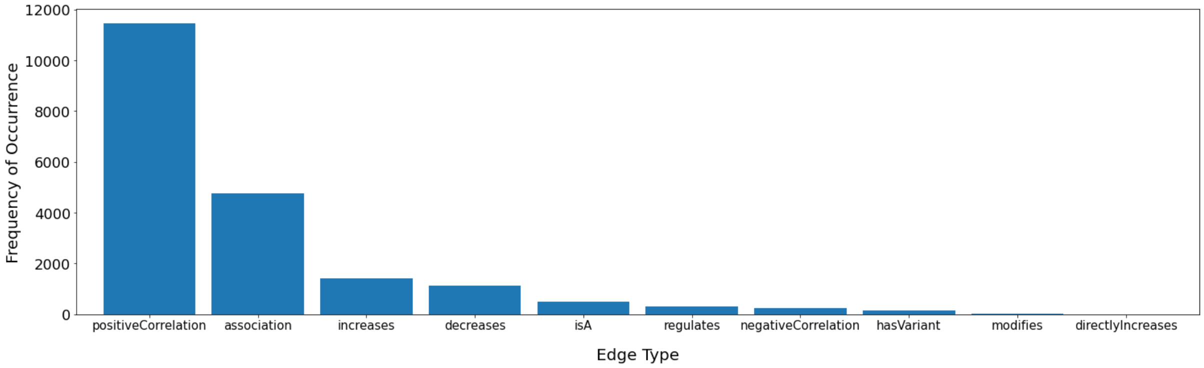


**Supplementary Figure 2. Edge type statistics of the schizophrenia knowledge graph.** Bar plot that represents the number of different BEL relations.

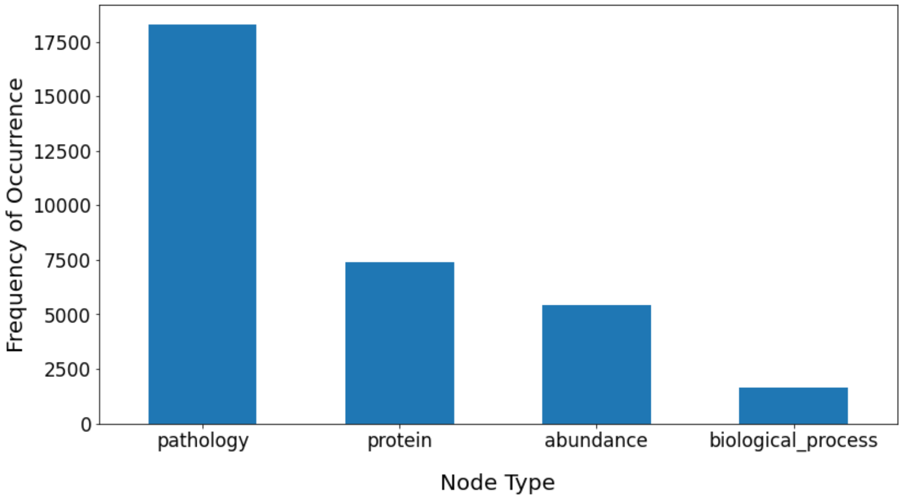


**Supplementary Figure 3. Node type statistics of the bipolar disorder knowledge graph. (a)** Bar plot representing the number of BEL statements that contain the different node/entity types.

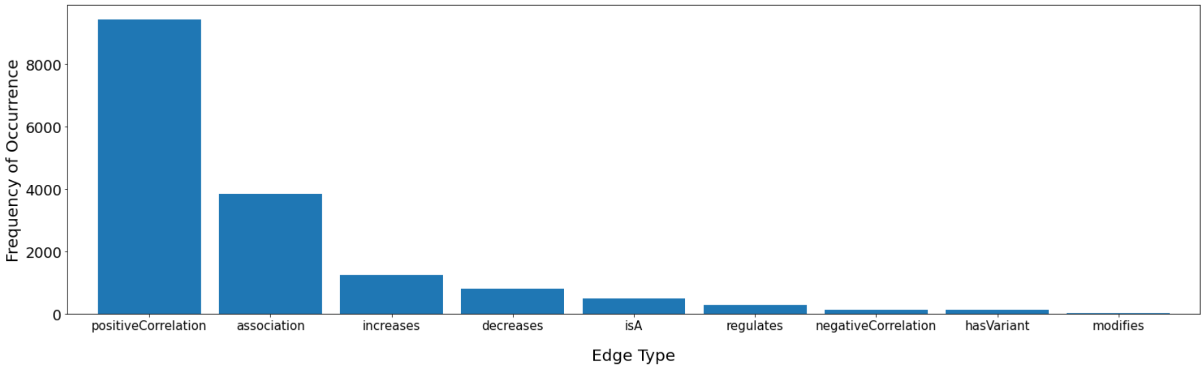


**Supplementary Figure 4. Edge type statistics of the bipolar disorder knowledge graph.** Bar plot that represents the number of different BEL relations.

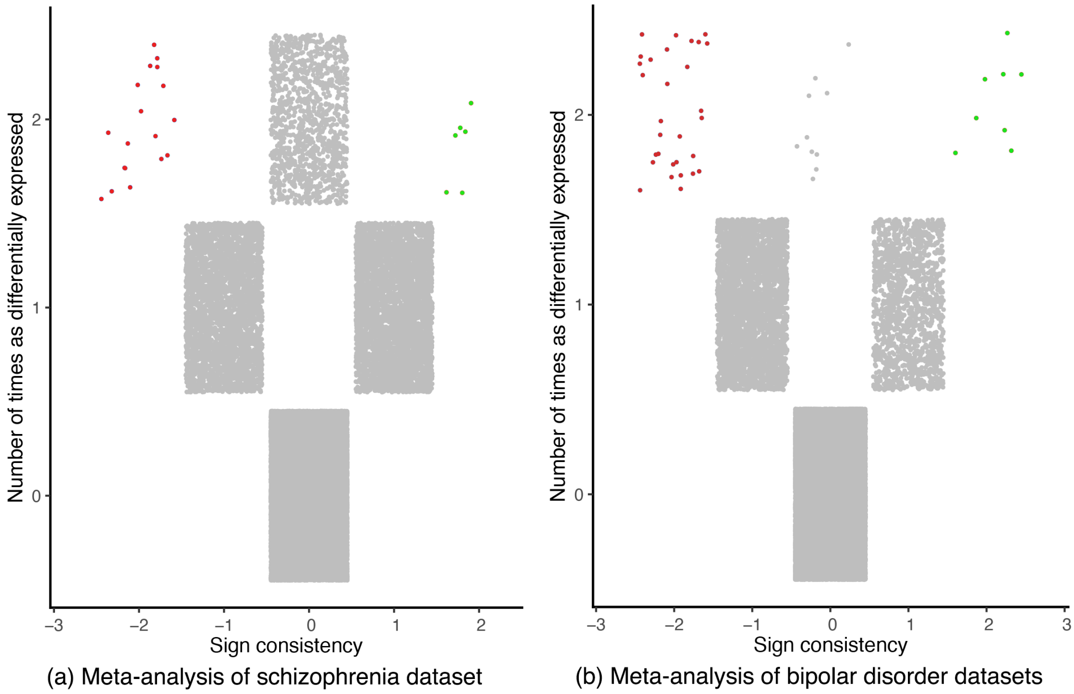


**Supplementary Figure 5. Meta-analysis results for the schizophrenia and bipolar disorder datasets.** Each scatter plot depicts the consistency in the direction of dysregulation (i.e., up- or down- regulated) of genes across datasets for **(a)** schizophrenia and **(b)** bipolar disorder (up-regulated genes in two or more datasets are highlighted in green and down-regulated genes in two or more datasets are highlighted in red.). The x-axis shows the sign of dysregulation (i.e., positive or negative) of a gene, and the total number of datasets that that direction is observed in. The y-axis provides the total number of datasets that the gene occurs in.

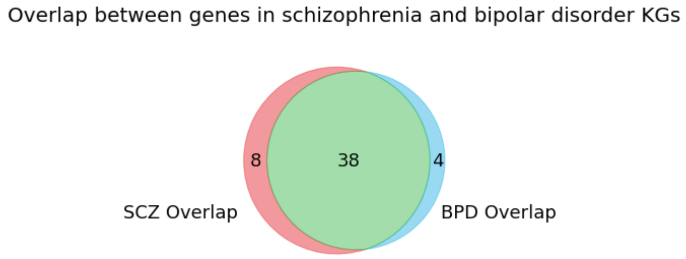


**Supplementary Figure 6. Overlap between genes in schizophrenia and bipolar disease KGs.** The figure shows the high overlap between genetically associated genes from GWAS Catalog present in each of the disease-specific KGs.
